# Hexanoic acid improves metabolic health in mice fed high-fat diet

**DOI:** 10.1101/2025.02.20.639216

**Authors:** Takako Ikeda, Yuki Nishimoto, Daisuke Ichikawa, Tomoka Matsunaga, Ami Kawauchi, Ikuo Kimura

## Abstract

**Background:** Overweight and obesity is currently a worldwide problem with undesirable health consequences such as type 2 diabetes. Therefore, much attention has been paid to preventing obesity through diet. Free fatty acids (FFAs) are an important energy source, and they also serve as signaling molecules in many biological processes leading to an increased energy expenditure and insulin secretion. Short-chain fatty acids (SCFAs) such as acetic, propionic and butyric acid are the bioactive metabolites produced by gut microbes, and their beneficial effects on host metabolism are well-studied. In addition, medium-chain fatty acids (MCFAs) such as octanoic and decanoic acid also play a positive role in regulating lipid and glucose metabolisms. However, the effects of hexanoic acid on metabolism are poorly understood. Therefore, this study investigated the role of hexanoic acid on lipid and glucose metabolism in mice.

**Methods:** Male C57BL/6J mice were fed normal chow diet, high-fat diet (HFD), HFD-containing 5% butyric acid or HFD-containing 5% hexanoic acid for 4 weeks, and the effects of hexanoic acid on lipid and glucose metabolism were examined.

**Results:** Butyric acid and hexanoic acid prevented body weight gain and fat accumulation in white adipose tissues under HFD-feeding. In addition, both FFAs suppressed the elevated plasma levels of non-esterified fatty acid (NEFA) and hepatic triglyceride content induced by HFD. The expression levels of genes involved in fatty acid biosynthesis were decreased in white adipose tissues by oral supplementation of butyric acid or hexanoic acid. Mice fed HFD also exhibited hyperglycemia and hyperinsulinemia, and these impaired glucose metabolisms improved by hexanoic acid. Hexanoic acid increased the expression levels of genes associated with gluconeogenesis, and improved insulin sensitivity in mice fed HFD.

**Conclusions:** This study highlights the importance of hexanoic acid in improvement of lipid and glucose metabolisms. Thus, our findings provide insight into the development of functional foods which prevent obesity-related disease such as type 2 diabetes.

## Introduction

It is currently estimated that more than a billion people are obese, and thus obesity is one of the major public health problems in the world (1). Overweight and obesity are associated with an increasing risk of several diseases such as type 2 diabetes, cardiovascular disease and cancers (2). Therefore, there is a growing need for the development and availability of functional foods with anti-obesity properties. As one of the most effective functional food ingredients, free-fatty acids (FFAs) have gained attention. FFAs are classified into the following 3 groups depending on their carbon chain length: short-chain fatty acids (SCFAs; less than 6 carbons), medium-chain fatty acids (MCFAs; 6-12 carbons) and long-chain fatty acids (LCFAs; more than 12 carbons). The majority of SCFAs in host are derived from microbial fermentation of indigestible carbohydrates, and MCFAs and LCFAs are mainly derived from dietary fat (3). SCFAs are microbial bioactive metabolites which are mainly composed of acetic (C2:0), propionic (C3:0) and butyric acid (C4:0) (4). SCFAs act as a mediator of the link between diet and metabolism, and their anti-obesity properties are well-studied (5). In particular, butyric acid exerts potent effects on metabolism. For example, butyric acid prevents diet-induced obesity and insulin resistance. In addition, butyric acid increases the release of a gut hormone glucagon-like peptide 1 (GLP-1), leading to an increased insulin secretion and a decreased food intake (6). MCFAs including hexanoic acid (C6:0) are abundantly in milk and coconut oil, and can be readily absorbed into the liver via the portal vein. In the liver, MCFAs are metabolized and used as fuel, indicating that absorption and metabolism of MCFAs are more rapid than that of LCFAs. Owing to their lipid properties, MCFAs have attracted much attention as an appropriate dietary oil for individuals with obesity and high energy demands (7). Accumulating evidence has demonstrated that MCFAs, especially octanoic acid (C8:0) and decanoic acid (C10:0), play an important role in maintaining glucose homeostasis and lipid metabolism. For example, oral administration of decanoic acid or medium-chain triglyceride (C10:0 triglyceride) protected from HFD-induced obesity and enhanced glucose tolerance through an increased GLP-1 secretion (8). In human studies, patients fed MCT diet showed enhanced oxygen consumption and thermogenesis, indicating that both SCFAs and MCFAs are potent modulators of lipid and glucose metabolism (9). However, the effects of hexanoic acid on metabolisms have largely remained elusive.

Hexanoic acid is a free-fatty acid with a 6-carbon chain length and commonly classified as a MCFA, but occasionally as a SCFA. This is because hexanoic acid can be taken from foods or produced by some anaerobic bacteria (10). In addition, hexanoic acid activates SCFA receptors such as GPR41 and GPR43, thereby showing unique properties (11). Only a few studies have investigated the effect of hexanoic acid on lipid and glucose metabolism. In human hepatoma HepG2 cell line, hexanoic acid inhibited the expression and activity of fatty acid synthase (Fasn), a key enzyme in de novo lipogenesis, induced by insulin and triiodothyronine (12). In LDL receptor knockout Leiden mice, which exhibit hyperinsulinemia, obesity and liver fibrosis after long-term HFD feeding, hexanoic acid did not attenuate the HFD-induced metabolic risk factors such as increased body weight, blood glucose, serum triglycerides (13). Therefore, the effects of hexanoic acid on metabolisms at whole-body level remain largely unknown. Herein, we examined whether hexanoic acid affects lipid and glucose metabolism using an HFD-induced obese mouse model.

## Materials and Methods

### Animal experiments

Male C57BL/6J mice were purchased from Japan SLC and housed at a temperature of 24 °C and 50% relative humidity under a 12 h light/dark cycle. The 6-week-old male mice were acclimated to the CLEA Rodent Diet (CE-2; CLEA Japan, Inc.) for 1 week, and randomly divided into 4 groups (n=10 for each group): normal chow diet (CE-2), HFD (D12492: 60% calories from fat; Research diets), HFD-containing 5% butyric acid (sodium butyrate; C4:0, FUJIFILM Wako), and HFD-containing 5% hexanoic acid (sodium hexanoate; C6:0, Tokyo Kasei). Mice were sacrificed under deep isoflurane-induced anaesthesia, and collected samples at 4 weeks after each dietary intervention. The body weight was measured once a week, and *ad libitum* food intake was assessed by calculating the average of daily food intake (g/day per mouse) for 4 weeks. Blood was collected from the inferior vena cava using heparinised injections, and plasma was separated by immediate centrifugation (7000 × g, 5 min, 4 °C). The compositions of the diets are shown in Supplementary Table S1. All experimental procedures were performed in accordance with the guidelines of the Committee in the Ethics of Animal Experiments of Kyoto University Animal Experimental Committee (Lif-K24002). All efforts were made to minimize suffering.

### Quantification of hexanoic acid

Hexanoic acid in the liver were determined according to a previous method (14). Briefly, the samples containing internal control (C19:0) were homogenized in methanol, followed by mixing with chloroform and water for lipid extraction. After centrifugation (2000 × g at 17 °C for 10 min), the supernatants were collected and dried. The samples resuspended in chloroform:methanol (1:3, v/v) were subjected to liquid chromatography with tandem mass spectrometry (LC-MS/MS) using an ultra-performance LC system (UPLC, Waters) equipped with an Acquity UPLC system coupled to a Waters Xevo TQD mass spectrometer (Waters). Using a methanol gradient in 10 mM ammonium formate aqueous solution, the samples were separated on an ACQUITY UPLC BEH C18 column (2.1 × 150 mm, 1.7 μm; Waters). The quantification of hexanoic acid was performed by using calibration curves.

### Biochemical analyses

Blood glucose level was measured using One Touch Ultra Test Strips (OneTouch® Ultra®; LifeScan). Plasma total cholesterol (LabAssay™ Cholesterol; FUJIFILM Wako), non-esterified fatty acid (NEFA) (LabAssay™ NEFA; FUJIFILM Wako), insulin (Insulin enzyme-linked immunosorbent assay (ELISA) kit (RTU); Shibayagi) levels were measured according to the manufacturer’s instructions. Hepatic triglyceride content was measured as previously described (15). Briefly, mouse liver was homogenized in a mixture of chloroform/methanol/0.45 M acetic acid, and the homogenate was rotated overnight at 4 °C. After centrifugation at 1500 × g for 10 min, the organic layer was collected, dried and resuspended in isopropanol. Plasma and hepatic triglyceride (LabAssay™ Triglyceride; FUJIFILM Wako) levels were measured according to the manufacturer’s instructions.

### Quantitative RT-PCR

Total RNA was isolated using an RNAiso Plus reagent (TAKARA). Reverse transcription was performed using Moloney murine leukemia virus reverse transcriptase (Invitrogen). cDNA was subjected to quantitative PCR analysis using the StepOne real-time PCR system (Applied Biosystems) with SYBR Premix Ex Taq II (TAKARA). Each value was normalized to 18S rRNA and calculated using the 2-ΔΔCt method. The primer sequences are shown in Supplementary Table S2.

### GTT and ITT

For the glucose tolerance test (GTT), mice fasted for 16 h were administered hexanoic acid (sodium hexanoate; C6:0, 2.5g/kg body weight, Tokyo Kasei) in 0.5% carboxymethylcellulose (CMC) by oral gavage. After 1 hr, mice were given glucose (1 g/kg body weight) by intraperitoneal injection. For the insulin tolerance test (ITT), mice fasted for 3 h were administered hexanoic acid 1 hr before intraperitoneally injection of insulin (0.75 mU/g; Sigma). In both tests, blood glucose levels were monitored at 0, 15, 30, 60, 90 and 120 min after glucose or insulin injection. 0.5% CMC was used as control.

### Statistical analysis

All data are presented as the mean ± SEM. The statistical analysis was performed using the GraphPad Prism (GraphPad Software Inc.). The Shapiro-Wilk test was used for the assessment of data normality. The statistical significance of differences between two groups was assessed by using a two-tailed unpaired Student’s *t*-test, whereas that of differences between multiple groups was assessed by using one-way ANOVA followed by the Tukey-Kramer post hoc test or Dunn post hoc test, depending on data normality. Statistical significance was set at P < 0.05.

## Results

### Oral administration of butyric acid or hexanoic acid prevents HFD-induced obesity

Previously, we showed that dietary SCFAs effectively suppressed HFD-induced obesity and improved metabolic functions (15). We thereby used butyric acid as a positive control, and examined whether hexanoic acid affects HFD-induced obesity. Male C57BL/6J mice were fed normal chow diet (ND), HFD, HFD-containing 5% butyric acid (HFD_C4) or HFD-containing 5% hexanoic acid (HFD_C6) for 4 weeks (Supplementary Table S1), and the hepatic level of hexanoic acid was measured. Feeding of HFD_C6 diet significantly increased the hepatic level of hexanoic acid, indicating that hexanoic acid was absorbed and directly transported to the liver by oral intake (Figure 1A). To examine whether hexanoic acid affects HFD-induced obesity, we measured the body weight. After 4 weeks of feeding, the body weights of mice fed HFD with hexanoic acid as well as with butyric acid were markedly decreased compared to those of mice fed HFD (Figure 1B). Although both sodium butyrate and sodium hexanoate possess the unpleasant flavor and odor, the amount of food intake did not change among all groups (Figure 1C). Therefore, these results indicate that hexanoic acid prevents HFD-induced obesity without reduced food intake, and the anti-obesity property of hexanoic acid is comparable to that of butyric acid.

**Figure 1.**
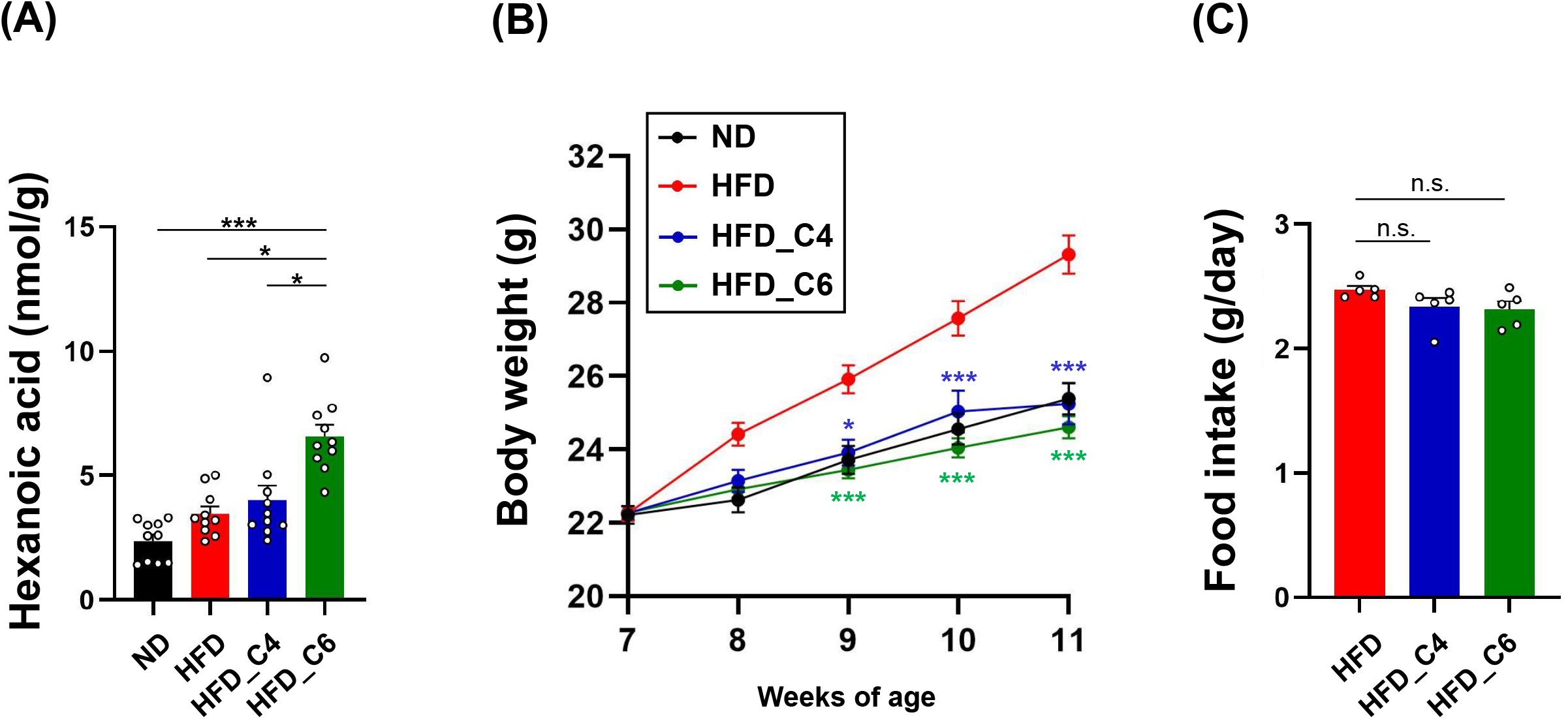
Oral supplementation of butyric acid or hexanoic acid improves HFD-induced obesity. **(A)** The liver content of hexanoic acid in mice fed normal chow diet (ND), high-fat diet (HFD), HFD-containing butyric acid (HFD_C4) or HFD-containing hexanoic acid (HFD_C6) (ND-fed group, n=10; HFD-fed group, n=10; HFD_C4 group, n=10; HFD_C6 group, n=10). **(B)** Body weight changes in mice fed ND, HFD, HFD_C4 or HFD_C6 (ND-fed group, n=10; HFD-fed group, n=10; HFD_C4 group, n=10; HFD_C6 group, n=10). **(C)** Food intake in mice fed HFD, HFD_C4 or HFD_C6 (HFD-fed group, n=5; HFD_C4 group, n=5; HFD_C6 group, n=5). All data are presented as the mean ± standard error of mean (SEM). ***P < 0.001; *P < 0.05, one-way ANOVA followed by the Dunn’s test (A) or the Tukey-Kramer test (B, D); Student’s *t*-test test, compared to HFD-fed group (C).

### Butyric acid and hexanoic acid decrease fat accumulation in mice fed HFD

We next performed the biochemical analyses to examine the effects of each diet on plasma lipids. In accordance to the previous report, the levels of plasma triglyceride (TG) were not changed among all groups, and the elevated levels of plasma total cholesterol in a group fed HFD were not also changed in groups fed HFD supplemented with butyric acid or hexanoic acid (Figure 2A, B) (15). However, the increased levels of plasma non-esterified fatty acids (NEFAs) induced by HFD-feeding were decreased by oral intake of butyric acid or hexanoic acid (Figure 2C). Obesity is primarily characterized by an excessive growth of adipose tissue mass. Even though mice were fed HFD only 4 weeks, HFD dramatically increased white adipose tissue mass (subcutaneous, perirenal, epididymal and mesenteric adipose). HFD-induced fat mass gain was suppressed in mice fed HFD containing either butyric acid or hexanoic acid to the same level as in the control mice fed ND (Figure 3A). Therefore, these results indicate that butyric acid and hexanoic acid possess potent anti-obesity properties by preventing fat accumulation in white adipose tissues. We then examined the mRNA expression levels of genes involved in lipogenesis and fatty acid oxidation in epididymal white adipose tissue. The mRNA expression levels of *Chrebp* and *Fasn*, which play an important role in the fatty acid biosynthesis, were up-regulated in mice fed HFD, and this increased expression of both genes was suppressed in mice fed HFD containing butyric acid or hexanoic acid (Figure 3B). In contrast, oral administration of butyric acid or hexanoic acid did not change the HFD-induced up-regulation of *PPARα* mRNA, which is involved in the fatty acid oxidation (Figure 3B). Hence, these results suggest that butyric acid and hexanoic acid exert anti-obesity effects through inhibiting lipid accumulation, rather than promoting energy expenditure in white adipose tissues.

**Figure 2.**
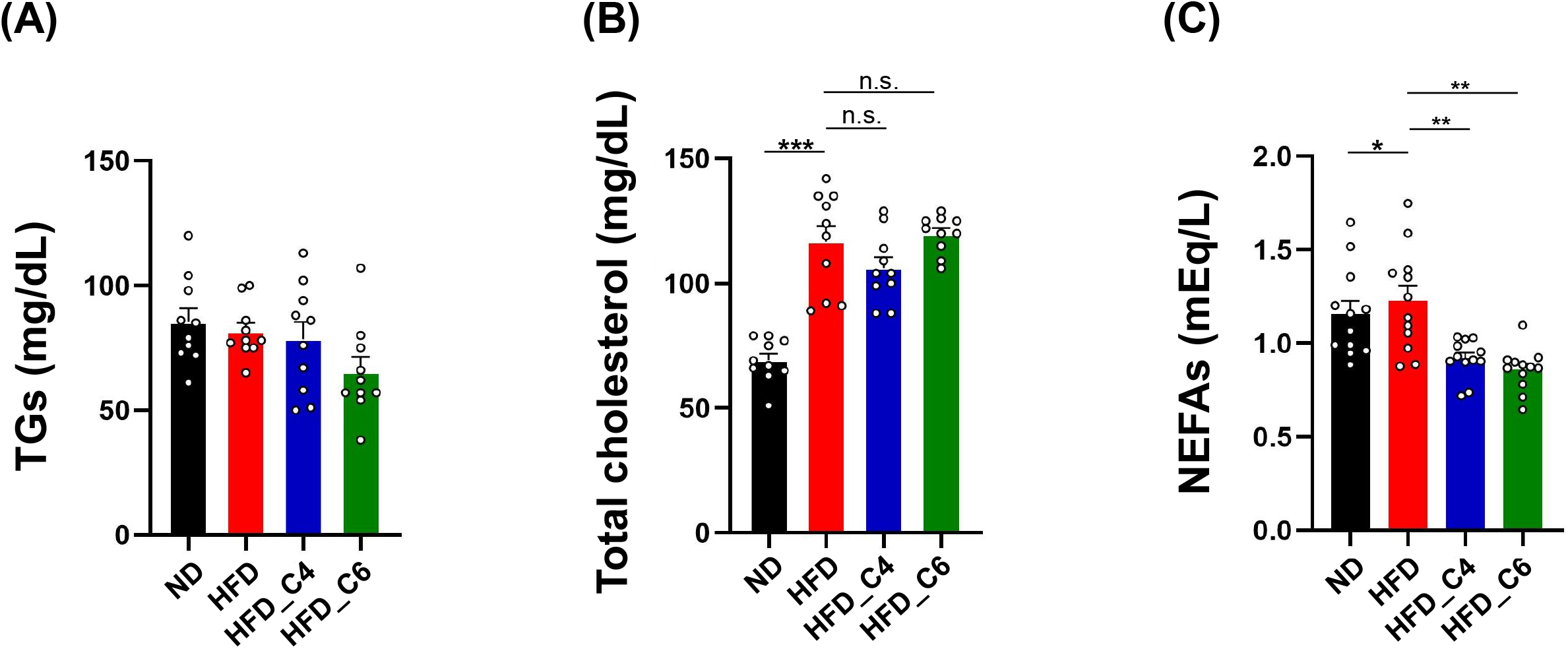
Biochemical analyses in mice fed HFD with or without butyric acid or hexanoic acid. **(A)** Plasma triglycerides (TGs), **(B)** plasma total cholesterol, **(C)** plasma non-esterified fatty acids (NEFAs) (ND-fed group, n=10; HFD-fed group, n=10; HFD_C4 group, n=10; HFD_C6 group, n=10). All data are presented as the mean ±SEM. ***P < 0.001; **P < 0.01; *P < 0.05; n.s., not significant, one-way ANOVA followed by the Tukey-Kramer test.

**Figure 3.**
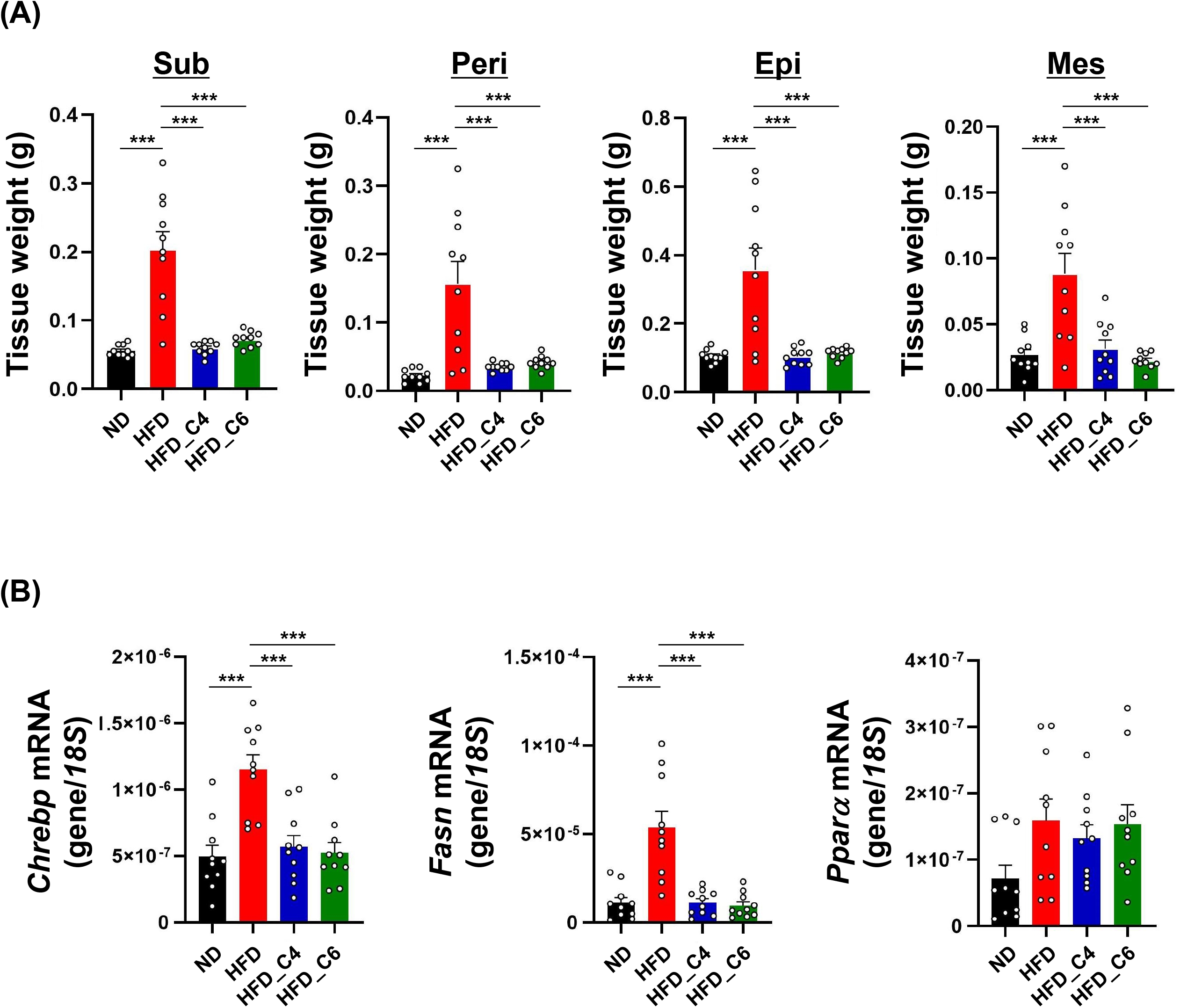
Butyric acid and hexanoic acid suppress lipid accumulation in adipose tissues under HFD feeding. **(A)** White adipose tissue weights in mice fed ND, HFD, HFD_C4 or HFD_C6 (ND-fed group, n=10; HFD-fed group, n=10; HFD_C4 group, n=10; HFD_C6 group, n=10). Sub, subcutaneous adipose tissue; peri, perirenal adipose tissue; epi, epididymal adipose tissue; mes, mesenteric adipose tissue. **(B)** The mRNA expression levels of genes involved in lipogenesis and fatty acid oxidation in epididymal adipose tissues. *Chrebp*: MLX interacting protein-like, *Fasn*: fatty acid synthase, *PPARa*: peroxisome proliferative activated receptor alpha. All data are presented as the mean ±SEM. ***P < 0.001, one-way ANOVA followed by the Tukey-Kramer test.

### Butyric acid and hexanoic acid decrease the hepatic TG contents in mice fed HFD

In contrast to LCFAs that are incorporated into chylomicrons as a triglyceride, dietary MCFAs are directly transported to the liver and rapidly metabolized by mitochondrial β-oxidation. Therefore, dietary MCFAs affect the hepatic lipid metabolisms. Indeed, octanoic acid (C8:0) and decanoic acid (C10:0) reduce hepatic lipid accumulation through a decreased lipogenesis and an increased lipolysis (16). We thus investigated the role of butyric acid and hexanoic acid in hepatic lipid metabolisms. Butyric acid and hexanoic acid effectively decreased the liver weight, and markedly reduced a higher content of hepatic TG in mice fed HFD (Figure 4A, B). However, the mRNA expression levels of *Chrebp, Fasn* and *PPARα* were not changed among each group (Figure 4C). The present results indicate that anti-obesity property of hexanoic acid, which is comparable with that of butyric acid, mainly exerts through inhibiting lipid accumulation in adipose tissues rather than through promoting β-oxidation in liver.

**Figure 4.**
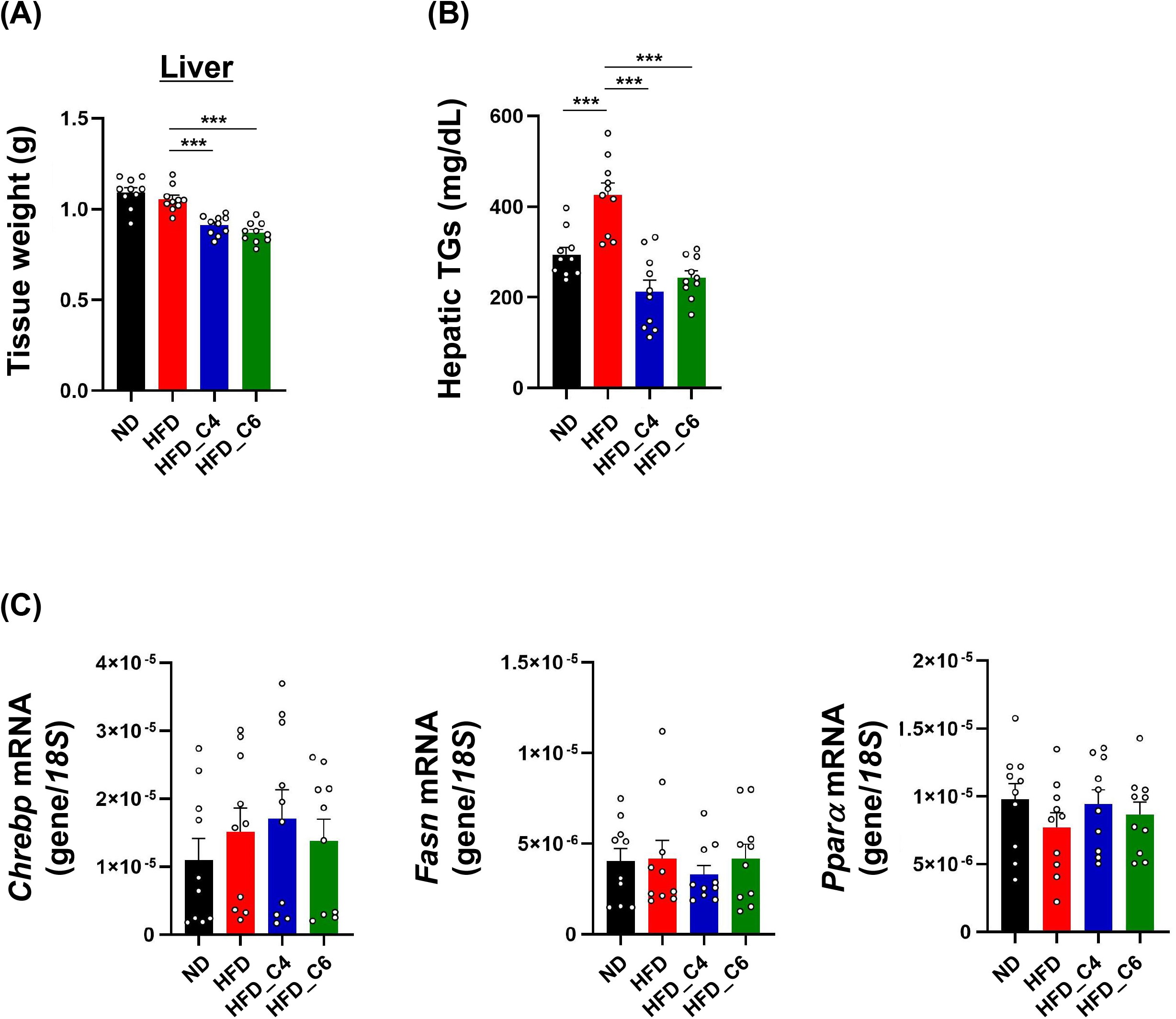
Butyric acid and hexanoic acid reduce the hepatic TG contents under HFD feeding. **(A)** Liver tissue weights and **(B)** hepatic TG contents in mice fed ND, HFD, HFD_C4 or HFD_C6 (ND-fed group, n=10; HFD-fed group, n=10; HFD_C4 group, n=10; HFD_C6 group, n=10). **(C)** The mRNA expression levels of genes involved in lipogenesis and fatty acid oxidation in liver. All data are presented as the mean ±SEM. One-way ANOVA followed by the Tukey-Kramer test or the Dunn’s test.

### Hexanoic acid improves glucose metabolism under HFD-feeding

Finally, we examined the effect of hexanoic acid on glucose metabolism. The elevated hepatic lipid deposition under HFD-feeding causes insulin resistance, and butyric acid protected from HFD-induced hyperinsulinemia and improved glucose tolerance (15, 17). We thus examined the blood glucose levels and found that hexanoic acid but not butyric acid significantly reduced the blood glucose under HFD-feeding (Figure 5A). In addition, both FFAs improved HFD-induced hyperinsulinemia, although butyric acid did not affect the HFD-induced hyperglycemia (Figure 5B). The mRNA expression levels of *Pepck* and *G6Pase*, which are associated with gluconeogenesis in the liver, were down-regulated in mice fed HFD (Figure 5C). Interestingly, HFD-induced decreased expression of both genes was restored by intake of hexanoic acid, indicating that hexanoic acid is more potent in maintaining glucose homeostasis than butyric acid (Figure 5C). We next performed intraperitoneal glucose tolerance test (IPGTT) and insulin tolerance test (ITT), and revealed that oral administration of hexanoic acid significantly enhanced glucose tolerance and insulin sensitivity (Figure 5D, E). Taken together, these results suggest that hexanoic acid is potent FFA which improves hyperglycemia under HFD-feeding through enhancing insulin sensitivity.

**Figure 5.**
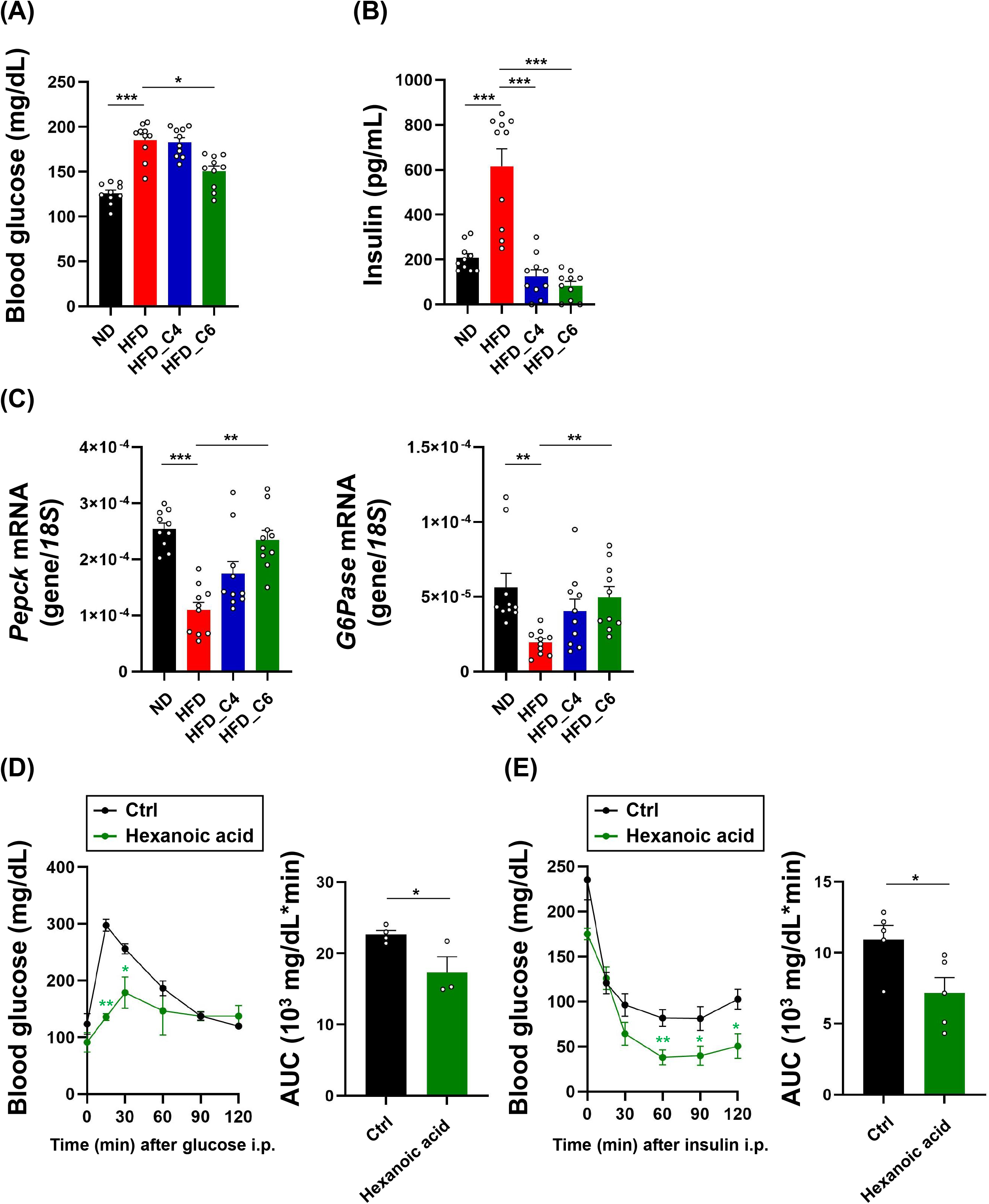
Hexanoic acid ameliorates hyperglycemia and hyper insulinemia in mice fed HFD. **(A)**Blood glucose and **(B)** plasma insulin levels in mice fed ND, HFD, HFD_C4 or HFD_C6 (ND-fed group, n=10; HFD-fed group, n=10; HFD_C4 group, n=10; HFD_C6 group, n=10). **(C)** The mRNA expression levels of genes involved in gluconeogenesis in liver. *Pepck*: phosphoenolpyruvate carboxykinase 1, *G6Pase*: glucose-6-phosphatase **(D)** Glucose tolerance test and **(E)** insulin tolerance test (Ctrl group, n=4; hexanoic acid group, n=3). All data are presented as the mean ±SEM. ***P < 0.001; **P < 0.01; *P < 0.05, one-way ANOVA followed by the Tukey-Kramer test (A, B) or the Dunn’s test (C); Student’s *t*-test test, compared to HFD-fed group (D).

## Discussion

In this study we investigated the effect of hexanoic acid on lipid and glucose metabolisms under HFD conditions, and found anti-obesity efficacy of hexanoic acid comparable to that of butyric acid (Figure 1B). Accumulating evidence has revealed that butyric acid prevents diet-induced obesity and insulin resistance. Oral supplementation of butyric acid enhances fatty acid oxidation and energy expenditure through up-regulating the expression of peroxisome proliferator-activated receptor gamma coactivator-1a and uncoupling protein 1 in brown adipose tissue (18). One mechanism by which butyric acid improves lipid and glucose metabolisms is by activation of SCFA receptors. The role of SCFA receptors in energy metabolism has been well-studied, especially for GPR41 and GPR43. Butyric acid exerts its effects through activating GPR43 because GPR43 exhibits a stronger response to longer SCFAs than GPR41. In our previous report, butyric acid conferred metabolic benefits on mice fed HFD, but these beneficial effects of butyric acid were abolished in *Gpr43* gene knockout mice (15). In this study, we confirmed that butyric acid inhibited HFD-induced body weight gain, fat accumulation in white adipose tissues and increased hepatic TG content (Figure 1B, 3A, 4B). Interestingly, hexanoic acid also improved obesity and lipid metabolism in mice fed HFD to the same extent as butyric acid, indicating potent efficacy of hexanoic acid in the treatment of obesity and obesity-related diseases. In addition, butyric acid and hexanoic acid suppressed the increased expression of *Chrebp* and *Fasn* under HFD-feeding, but not of PPARα, in adipose tissues. It suggests that both FFAs exert their effects in part through by inhibiting lipogenesis rather than by promoting fatty acid oxidation (Figure 3B). Although underlying mechanisms linking hexanoic acid to a metabolic benefit remain unclear, our results suggest that hexanoic acid, as well as butyric acid, is a potent FFA with anti-obesity property.

Butyric acid also plays an important role in the regulation of glucose metabolism and insulin sensitivity. For example, butyric acid promotes incretin secretion and gluconeogenesis in the intestine via the gut-brain axis (6, 19). Besides, supplementation of inulin-type fructan, which promotes the growth and activity of butyric acid-producing bacteria, improves glucose intolerance (20). In this study, we showed that butyric acid improved the hyperinsulinemia of mice fed HFD, but failed to suppress hyperglycemia (Figure 5A, B). Consistent with these results, the decreased expression of *Pepck* and *G6Pase* in liver of mice fed HFD were not fully restored, but tended to exhibit a mild increase (Figure 5C, D). In contrast, hexanoic acid improved the hyperinsulinemia and reduced blood glucose levels (Figure 5A, B). Furthermore, hexanoic acid fully restored the decreased expression of *Pepck* and *G6Pase* in mice fed HFD, and increased insulin sensitivity (Figure 5C, D, E, F). Therefore, these results indicate that hexanoic acid is a remarkable FFA with anti-diabetes property.

SCFAs are the major end-products of microbial fermentation in both ruminants and non-ruminants, and their amounts and components of tissues differ markedly among species. In ruminant animals, SCFAs are produced from microorganisms in forestomach and rapidly absorbed, which provide nearly 80% of the energy requirement of the animal (21). In mammals including human, majority of SCFAs produced in the colon is utilized by epithelial colonic cells as fuel, and thus they provide only 10% of the energy required by cells in the whole body (22). Therefore, a higher amount of SCFAs and hexanoic acid is present in the peripheral circulation of ruminant animals than that of mammals. The level of circulating FFAs affects milk fat composition. Cow milk, for example, contains hexanoic acid, but human breast milk does not. It indicates that dietary intake of hexanoic acid is required for us to improve metabolic health. Obesity and obesity-related diseases are now worldwide health problems. To solve these problems, effective therapeutic strategies such as developing drugs and supplements are required. Although the molecular mechanisms behind the anti-obesity and anti-diabetic effects of hexanoic acid remain elusive, hexanoic acid is a promising candidate for the treatment of metabolic disorders. Taken together, our results help understanding the role of hexanoic acid in improving metabolic function.

## Data availability statement

All data are included in this article and its Supplementary material, and are available from the corresponding authors upon reasonable request.

## Ethics statement

All experimental procedures using mice were performed according to protocols approved by the Committee on the Ethics of Animal Experiments of the Kyoto University Animal Experimentation Committee (Lif-K24002).

## Author contributions

TI performed the experiments, interpreted data, and wrote the paper; NY, ID, MT and KA performed the experiments; IK supervised the project and interpreted data; TI and IK had primary responsibility for the final content. All authors read and approved the final manuscript.

## Acknowledgments

This work was supported by research grant from the Food Science Institute Foundation (Ryoshoku Kenkyukai, Japan).

