## Supplemental material for "Hexanoic acid improves metabolic health in mice fed high-fat diet"

**Supplementary Material**

**Supplementary Table1.** Dietary composition of high-fat diet used in this study. HFD, high-fat diet; HFD_C4, HFD containing 5% butyric acid; HFD_C6, HFD containing 5% hexanoic acid.

**Supplementary Table2.** Primer sequences used in this study.

| Formula | HFD | HFD_C4 | HFD_C6 |
|---|---|---|---|
| Product | | kcal % | |
| Protein | 20 | 20 | 20 |
| Carbohydrate | 20 | 20 | 20 |
| Fat | 60 | 60 | 60 |
| Ingredient | | gm | |
| Casein, 30 mesh | 200 | 190 | 190 |
| L-cystine | 3 | 2.85 | 2.85 |
| Maltodextrin 10 | 125 | 118.25 | 118.25 |
| Sucrose | 68.8 | 65.36 | 65.36 |
| Cellulose, BW200 | 50 | 47.5 | 47.5 |
| Butyric acid | 0 | 38.6925 | 0 |
| Hexanoic acid | 0 | 0 | 38.6925 |
| Soybean oil | 25 | 23.75 | 23.75 |
| Lard | 245 | 232.75 | 232.75 |
| Mineral mix S10026 | 10 | 9.5 | 9.5 |
| Dicalcium phosphate | 13 | 12.35 | 12.35 |
| Calcium carbonate | 5.5 | 5.225 | 5.225 |
| Potassium citrate. 1 $H_2O$ | 16.5 | 15.675 | 15.675 |
| Vitamin mix V10001 | 10 | 9.5 | 9.5 |
| Choline bitartrate | 2 | 1.9 | 1.9 |
| FD&C blue dye* | 0.05 | 0.0475 | 0.0475 |

*FD&C blue dye: synthetic organic compound primarily used as a blue colorant for dietary supplements.

**Supplementary Table S1**

| | Forward | Reverse |
|---|---|---|
| *18s* | 5'-acgctgagccagtcagtgta-3' | 5'-cttagagggacaagtggcg-3' |
| *Chrebp* | 5'-ctggggacctaaacaggagc-3' | 5'-gaagccaccctatagctccc-3' |
| *Fasn* | 5'-gctgcggaaacttcaggaaat-3' | 5'-agagacgtgtcactcctggactt-3' |
| *PPAR* $\alpha$ | 5'-cctgaacatcgagtgtcgaa-3' | 5'-ggccttgaccttgttcatgt-3' |
| *Pepck* | 5'-ccacagctgctgcagaaca-3' | 5'-gaagggtcgcatggcaaa-3' |
| *G6Pase* | 5'-ccatgcaaaggactaggaacaa-3' | 5'-taccagggccgatgtcaac-3' |

**Supplementary Table S2**
